# Deformable Models-Based Retinal OCT Layer Segmentation and Classification with Feature Analysis

**DOI:** 10.64898/2026.07.20.739618

**Authors:** Maria V. Leyba-Mesa, Bayan Ahmad, Elijah Ray, Aryan Patel, Buket D. Barkana

## Abstract

Optical coherence tomography (OCT) is widely used for retinal disease assessment, but automated quantitative analysis remains challenging because of anatomical variability and noisy imaging conditions. This study presents an interpretable OCT classification framework based on four anatomically guided retinal layers, combining preprocessing, adaptive segmentation, targeted feature engineering, and supervised classification to identify Normal, CNV, DME, and Drusen cases. Layer-specific descriptors included statistical, derivative, fluid-related, and GLCM texture markers. Feature correlation and ranking analyses showed that the proposed descriptors were highly complementary, that the most informative features were concentrated in layers 2 and 4, and that layers 1 and 3 contributed supportive structural information. Among the evaluated classifiers, the neural network performed best, achieving an accuracy of 98.17%, sensitivity of 97.88%, specificity of 99.38%, and AUC of 0.9985. Computational analysis showed efficient training and inference, with a total training time of 2216.3 s, prediction speed of approximately 160000 observations per second, and a compact model size of about 11 kB. These results demonstrated that anatomically guided feature extraction can provide accurate, efficient, and interpretable OCT disease classification, offering a practical alternative to less transparent end-to-end deep learning approaches.

## Introduction

Approximately 288 million people worldwide are projected to be affected by age-related macular degeneration (AMD) by the year 2040 (Kaur and Singh, 2023), and 161 million people by the year 2045 with diabetic macular edema (DME) (Wong and Tan, 2023). AMD and DME are two of the most prominent retinal diseases. The retina, the innermost layer of the eye, plays a major role in converting light signals into electrical signals (Mahabadi and Al Khalili, 2024). Thus, damage to retinal tissue causes a mistranslation between the two signal types and hinders the brain’s perception. This misperception directly correlates to the visual impairment experienced by those with retinal diseases. Retinal diseases are often assessed using optical coherence tomography (OCT), which allows ophthalmologists to view a cross-section of the retina (Fu et al., 2016) to determine disease type and progression and to advise on treatments. With the projected increase in AMD and DME cases, developing robust automated methods for retinal OCT image segmentation and analysis is becoming more important.

The retina comprises ten layers, some with vastly different contents and functions (Figure 1). The inner limiting membrane (ILM) is the innermost subretinal layer that separates the retina from the vitreous humor. This is followed by the nerve fiber layer (NFL). Ganglion cells are housed in the ganglion cell layer. A series of subsequent plexiform and nuclear layers comprises much of the retinal structure. The rod and cone cell bases are located in the external limiting membrane (ELM) (Mahabadi and Al Khalili, 2024), which also separates the preceding layers from the photoreceptor segments and the retinal pigment epithelium (RPE) (Fu et al., 2016). The posterior RPE also comprises some of the Bruch’s membrane complex (Borelli et al., 2018). The choroid i provides a blood supply to the outermost layers of the retina (Mahabadi and Al Khalili, 2024). The normal aging of the eye is reflected in OCT scans. However, these changes are presented differently than in AMD- and DME-conditioned eyes. In young, healthy retinas, OCT images show clearly defined, continuous alternating hyperreflective and hyporeflective bands, with thin and smooth outer retinal layers. With normal aging, the ELM and outer bands may thicken, become more irregular, and show reduced visibility and boundary definition (Chen et al., 2023).

**Figure 1.**
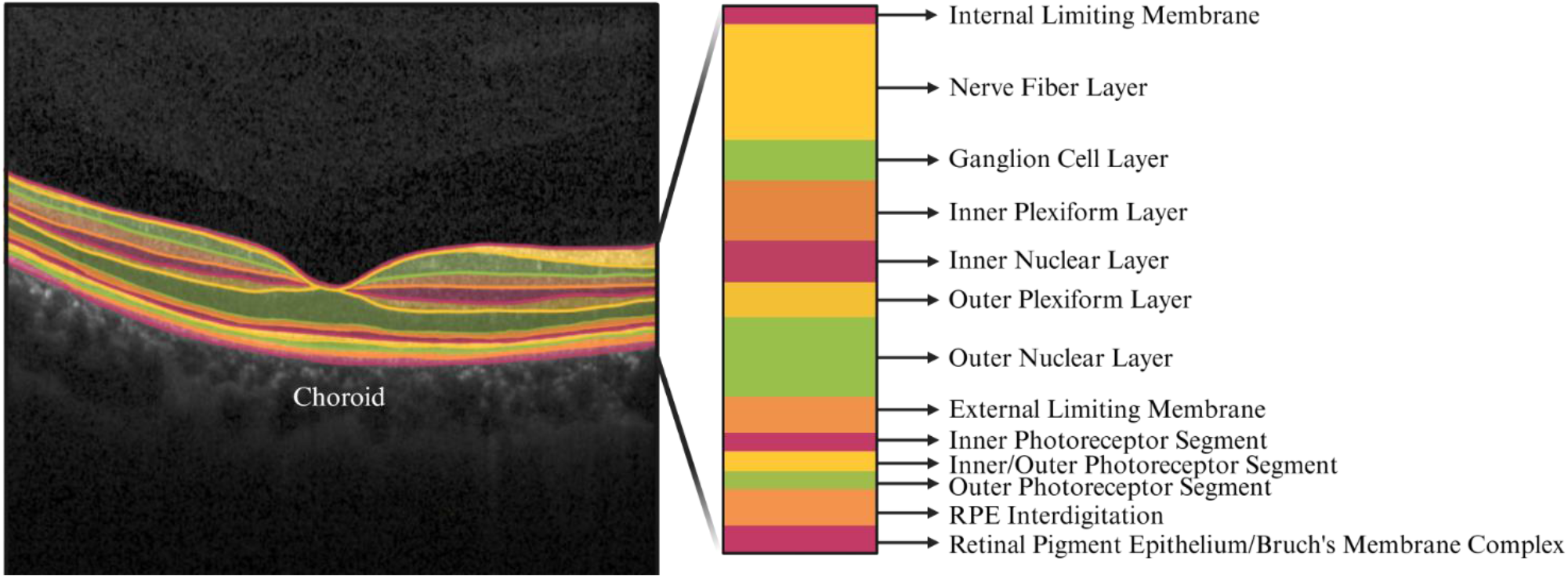
Retinal layers of a normal (healthy) case on an OCT image.

It is reported that AMD results from inflammatory responses to oxidative stress associated with aging (Hollyfield et al., 2008). It is associated with drusen depositions, which are aggregates of extracellular components. There are two branches of AMD: dry and wet. Dry AMD is more prevalent and characterized by a thicker accumulation of drusen, along with a more prominent Bruch’s membrane and a loss of photoreceptors. Choroidal neovascularization (CNV) is a critical feature of wet AMD and results from subretinal cell migration (Kaur and Singh, 2023). Diabetes mellitus causes diabetic Retinopathy (DR) and diabetic macular edema (DME), which presents in DR as fluid pockets that develop in the neural retina, causing it to thicken (Kaur and Singh, 2023).

### Previous works

Prior studies have shown that retinal layer thickness, hyperreflective retinal spots, disorganization of the retinal inner layers, outer retinal integrity, and related retinal OCT biomarkers can provide clinically meaningful information about disease status and prognosis (Vujosevic et al., 2023; Pandya et al., 2024, Bi et al., 2025, Zhang et al., 2025). Therefore, computational analysis of OCT images has increasingly shifted from simple image visualization toward objective measurement of retinal morphology.

A major step in retinal OCT-based quantitative analysis is retinal layer segmentation. Accurate segmentation is necessary before thickness, shape, reflectivity, and layer-specific structural features can be extracted. Earlier reviews emphasized that OCT segmentation remains challenging because OCT images are affected by speckle noise, low contrast, shadow artifacts, and disease-related disruptions of normal retinal anatomy (Kafieh et al., 2013). Traditional segmentation approaches have included A-scan- and B-scan-based methods, active contours, graph-based methods, edge detection, and artificial intelligence-based methods. Although many methods can perform well on limited or relatively normal datasets, segmentation robustness remains a challenge in pathological images, where retinal boundaries may become weak, distorted, or discontinuous.

Liu et al. (2022) developed an automated retinal boundary segmentation method using an improved Canny operator to reduce errors caused by vascular shadows, vitreous artifacts, noise, discontinuities, and false edges. The method segmented eleven retinal boundaries and was evaluated against ophthalmologist annotations. Average differences were approximately 2–6 µm in healthy eyes and 3–10 µm in age-related macular degeneration cases, with 98% of healthy and 94% of AMD segmentations rated “perfect” or “good.” Its main strengths are interpretability, strong boundary accuracy, and potential use as a standalone method or preprocessing stage. However, it addresses boundary detection rather than feature ranking or disease classification, and broader validation across multiple retinal disorders is needed to establish generalizability across diverse diseases and imaging devices.

Mittal and Bhatnagar (2023) proposed a retinal OCT segmentation framework that integrates speckle-noise reduction, weak-boundary detection, and multilayer delineation. Their approach combined a far-flung ratio algorithm, a random walk and inter-frame flattening methods, and N-ret layer segmentation. Reported performance included 97.25% accuracy, 98.258% sensitivity, 96.32% recall, 97% noise reduction, and 13.94 s execution time. A key strength was its direct treatment of speckle noise and low-intensity boundaries, outperforming Canny edge, two-pass edge, and edge-flow techniques. However, validation focused mainly on segmentation and denoising performance and did not include large, multi-device datasets or diverse retinal pathologies, thereby limiting evidence for broader clinical generalizability.

Sakthi Sree Devi et al. (2021) developed an OCT-based method for detecting diabetic retinopathy. The retina was segmented into seven layers using graph cuts, and layer thickness and neovascularization features were extracted from 8 normal and 5 diabetic retinopathy images. The study identified differences in retinal thickness, including increased nerve fiber layer thickness and reduced inner nuclear layer thickness in diabetic retinopathy, supporting the value of interpretable structural features. However, the very small dataset and the lack of feature ranking, cross-validation, classifier evaluation, and comparisons among supervised models limited generalizability and robustness.

Roy et al. (2025) proposed a graph-based method for segmenting spectral-domain retinal OCT images, using a shortest-path search to delineate seven intra-retinal boundaries and estimate six layer thicknesses. The study compared Gaussian and wavelet denoising and reported a substantial reduction in analysis time, with automated processing taking 4.93 s versus 578.05 s for manual segmentation. Wavelet denoising improved accuracy but added approximately 10 s per image, whereas Gaussian filtering provided faster, more practical preprocessing. Key strengths include validation against manual segmentation and explicit consideration of computational efficiency, both relevant to clinical implementation. However, the work focused on segmentation and thickness estimation and did not investigate feature ranking or supervised disease classification. In addition, the use of healthy adult retinas may limit the method’s demonstrated robustness and generalizability to pathological OCT images.

Hossain et al. (2025) proposed a 3D SD-OCT layer segmentation method that used gradient intensity, ROI selection, Canny edge detection, graph formulation, and Dijkstra’s shortest-path algorithm. By incorporating volumetric context, the approach aimed to improve retinal boundary detection beyond conventional 2D methods. Tested on 288 B-scans from 12 patients, it achieved mean RMSEs of 2.82, 4.88, 2.03, 3.77, and 0.64 pixels across five boundaries. Strengths included quantitative validation and comparison with recent methods; limitations were the small sample size and potentially greater computational complexity relative to simpler 2D segmentation pipelines on larger datasets.

Literature has established a strong foundation for retinal OCT segmentation and OCT-based disease assessment. However, many existing studies focus solely on segmentation performance, while others proceed directly to deep learning-based image classification. Relatively fewer studies examine how quantitative features derived from segmented retinal layers contribute to classification performance. In our study, we address this gap by using a stepwise deformable model-based pipeline that includes retinal layer segmentation, feature extraction, feature ranking, and supervised classification. It allows the classification results to remain associated with measurable retinal layer features rather than relying only on raw image-based prediction. This study makes the following contributions:

- It introduces an interpretable OCT classification framework based on four anatomically guided retinal layers, enabling examination of disease-related patterns within distinct structural regions.
- It develops a layer-specific feature representation that comprises statistical, derivative-based, fluid-related, and texture descriptors to characterize complementary retinal properties.
- It demonstrates through feature-ranking analysis that the most discriminative information is derived predominantly from layers 2 and 4.
- It evaluates supervised machine learning classifiers.
- It evaluates prediction speed and model size, demonstrating that the proposed framework is computationally efficient and well suited for rapid OCT image analysis.

The remainder of this paper is organized as follows. Materials and Methods presents the computational framework for OCT preprocessing, retinal layer segmentation, feature extraction and ranking, and neural-network classification. Experimental Results and Discussion evaluates segmentation performance, feature importance, and computational efficiency. The Conclusion summarizes findings and outlines future research directions.

## Materials and Methods

The proposed methodology comprises four main stages: retinal OCT image preprocessing, four-layer retinal segmentation, quantitative feature extraction, and supervised classification (Figure 2). First, each image is preprocessed to reduce noise and enhance the visibility of layer boundaries. A four-layer retinal segmentation method is then applied to partition the retina into anatomically meaningful regions for subsequent quantitative analysis. The first layer includes the inner limiting membrane and the nerve fiber layer; the second layer includes the tissue between the nerve fiber layer and the external limiting membrane; the third layer contains the photoreceptor segment and adjacent retinal regions; and the fourth layer represents the choroid. After segmentation, layer-specific quantitative descriptors are extracted and used as inputs to the classification stage. Figure 2 illustrates the complete workflow, showing how raw images are transformed into quantitative retinal features for computational disease classification.

**Figure 2.**
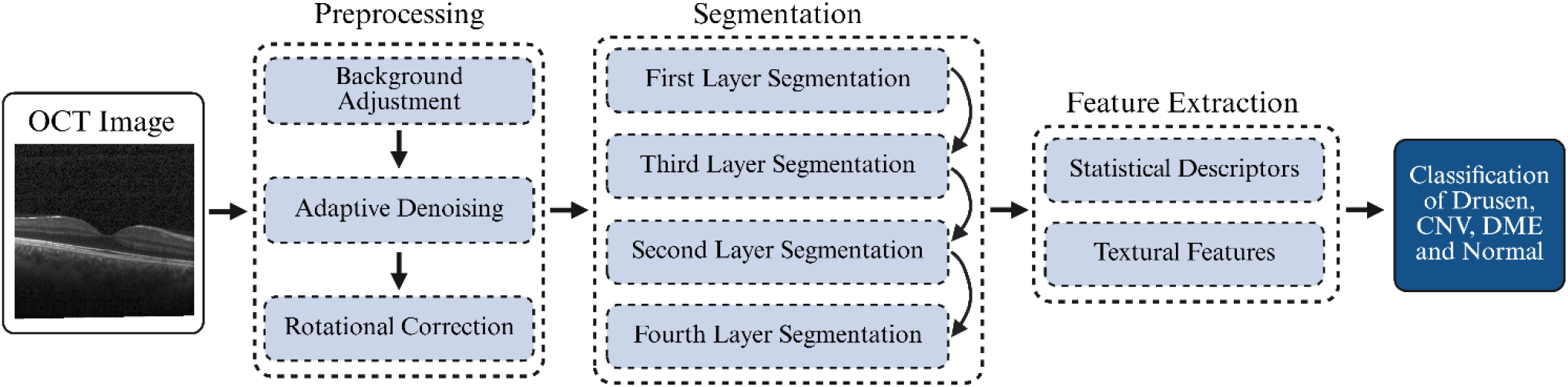
Block diagram of the proposed OCT image analysis pipeline, illustrating the sequential stages of preprocessing, retinal layer segmentation, feature extraction, and disease classification.

### Dataset

The dataset contains 84,495 retinal OCT images in JPEG format, organized into four diagnostic categories: NORMAL, CNV, DME, and DRUSEN (Kermany et al., 2018). The class distribution is 27,323 NORMAL images, 37,206 CNV images, 11,349 DME images, and 8,617 DRUSEN images. Representative images from the four categories are shown in Figure 3.

**Figure 3.**
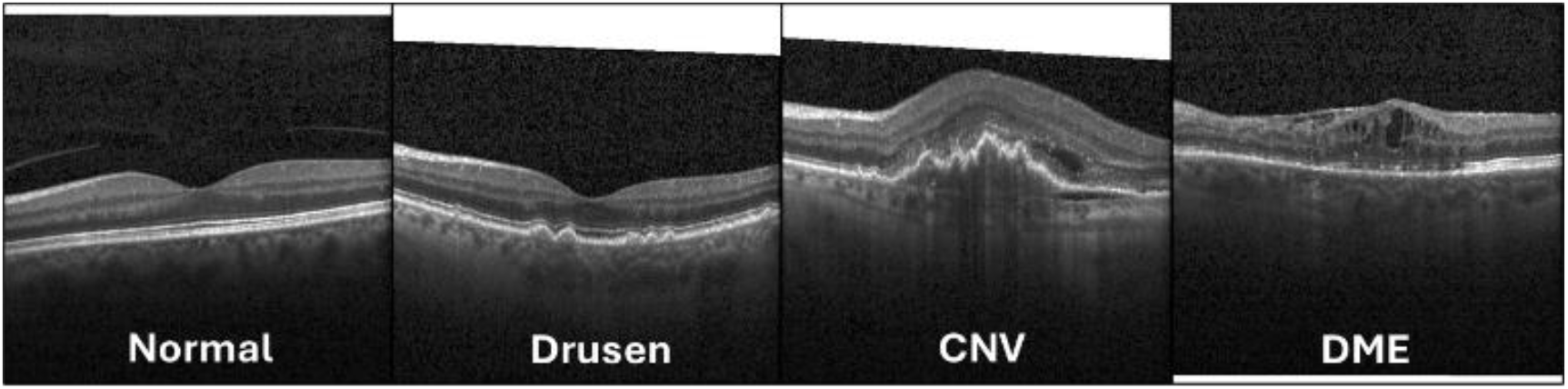
Representative OCT images from the four classes used in this study: Normal, Drusen, CNV, and DME, illustrating the characteristic structural differences observed across retinal disease categories (Leyba-Mesa et al., 2025).

### Background Adjustment

Some OCT images contained bright white regions along the outer edges of the retinal images. The Normal, Drusen, and CNV images in Figure 3 show white regions that could interfere with subsequent segmentation by introducing irrelevant background information. To reduce this effect, the preprocessing stage begins with a background masking step that isolates and removes these unwanted regions before further analysis. The implementation first inverts the input image, then scans its boundaries to detect the transition from the retinal region to the white background. The detected edge coordinates are used to define the valid image extent, and all pixels outside this region are set as a part of the background.

### Adaptive Denoising

The dataset contains images with a significantly low signal-to-noise ratio due to noise. We employed a histogram-based thresholding approach to filter noise, as illustrated in Figure 4a. An adaptive thresholding value was determined based on the processed image’s noise level. A *K*×*L* block of the background with distributed noise, *I*_*n*_, was used, where *K* and *L* were set to 50. The block’s average pixel intensity value, *µ*_*n*_, was used to define an adaptive threshold for filtering noise. Assuming the pixel intensity values of *I*_*n*_ have a relatively normal distribution, approximately half of the pixels in *I*_*n*_ should have an intensity value below *µ*_*n*_ due to the shape of the cumulative distribution curve and the position of *µ*_*n*_. For any pixel in *I*_*n*_ the univariate normal pdf (Stark and Woods, 1986) can be defined as

**Figure 4.**
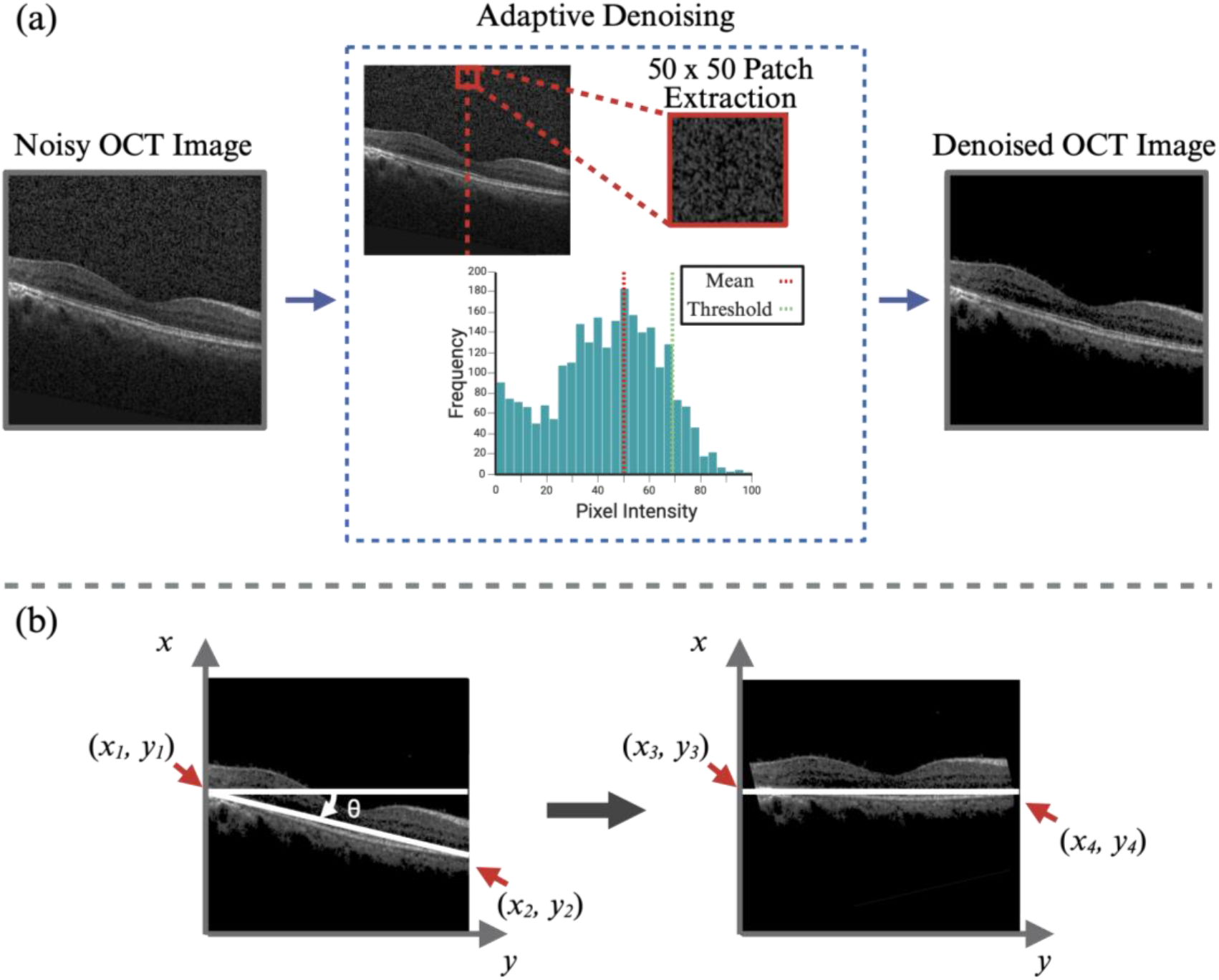
(a) Overview of the adaptive denoising pipeline for OCT images, showing a noisy input image, local intensity extraction from a 50×50 patch, histogram-based mean thresholding, and the resulting denoised OCT output. (b) Illustration of the rotational correction procedure for tilted OCT images showing the original slanted retinal boundary, the estimated rotation angle *θ*, and the corrected image after alignment (Leyba-Mesa et al., 2025).

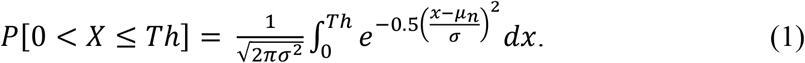

Any pixel intensity, *x*, obeys the normal probability law with mean *µ*_*n*_ and standard deviation *σ*. A threshold value, *Th*, was determined to be at the value of *µ*_*n*_ + *σ* to eliminate the background while keeping the integrity of the region of interest (ROI) edges. It corresponded to approximately 85% of the pixels within *I*_*n*_. Algorithm 1 summarizes the steps of the implemented adaptive denoising.

#### Algorithm 1

Adaptive Denoising Filter

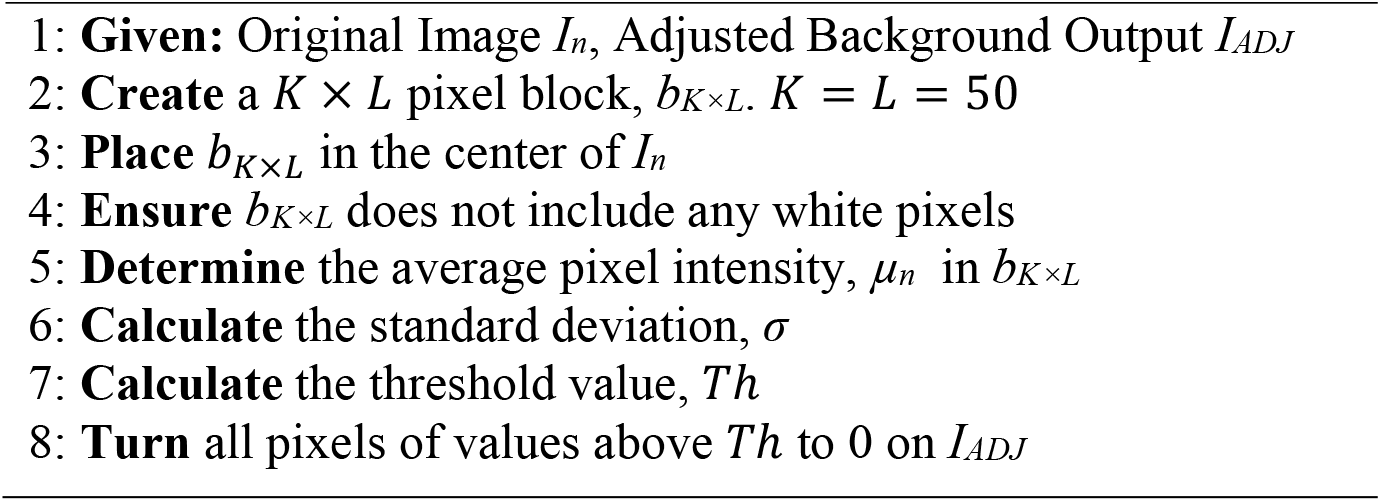

### Rotational Correction

Due to the nature of retinal OCT image acquisition, retinal layers appear tilted in some images, as shown in Figure 4b. Because the proposed layer segmentation model performs more reliably when the retinal layers are horizontally aligned, a custom image-rotation step was implemented before segmentation. This method automatically detects the tilt angle of the retinal layers and rotates the image to align the layers horizontally. By standardizing layer orientation, this preprocessing step improves the consistency and accuracy of the four-layer segmentation.

The rotation process first detects the coordinates of the photoreceptor segment; a simple thresholding method is sufficient since the segment has the highest image intensity. The rotation angle required to align the image was calculated using the arctangent function in (2). Δy is the vertical distance and Δx is the horizontal distance.

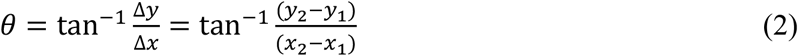

The angle *θ* was applied to align the detected edge points horizontally, thus correcting any tilt. The rotation ensures that all images are consistently aligned, minimizing errors caused by tilt and improving the accuracy of subsequent processing steps.

New coordinates, (*x*_3_, *y*_3_) and (*x*_4_, *y*_4_), are calculated using equations 3 and 4. Algorithm 2 summarizes the steps.

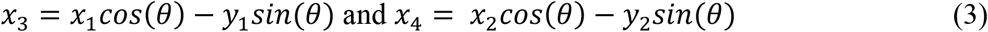

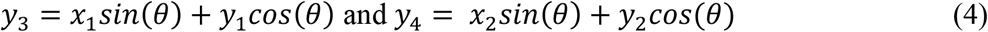

#### Algorithm 2

Rotational Correction

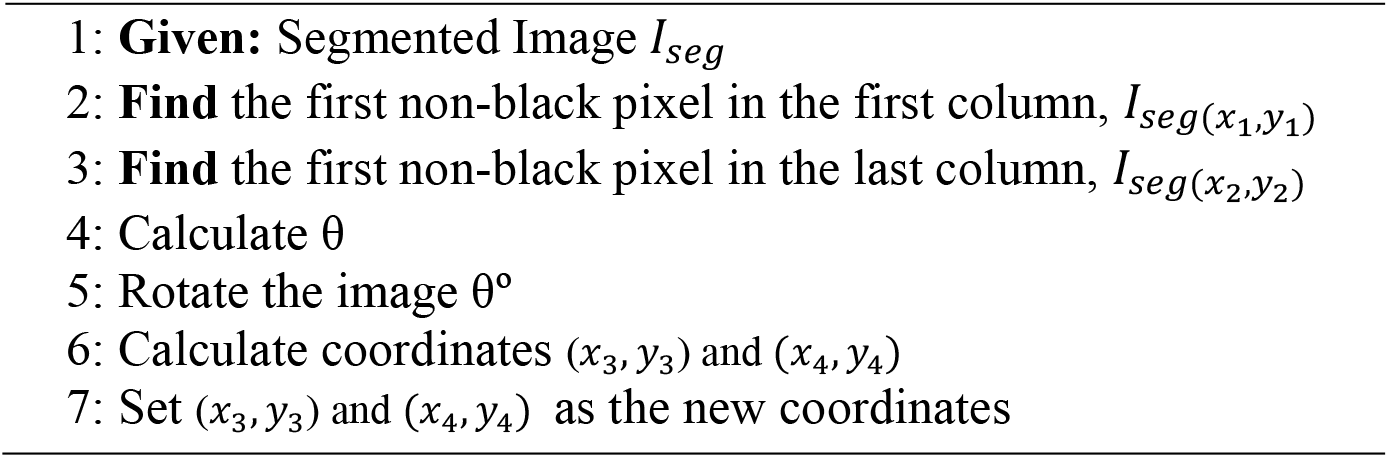

### Segmentation

Retinal layer segmentation was performed as an initial step to isolate the major structural regions of the OCT image for subsequent feature extraction and classification. The proposed approach uses a rule-based image-processing pipeline (Leyba-Mesa et al., 2025). The overall workflow of the proposed four-layer segmentation method is summarized in Figure 5.

**Figure 5.**
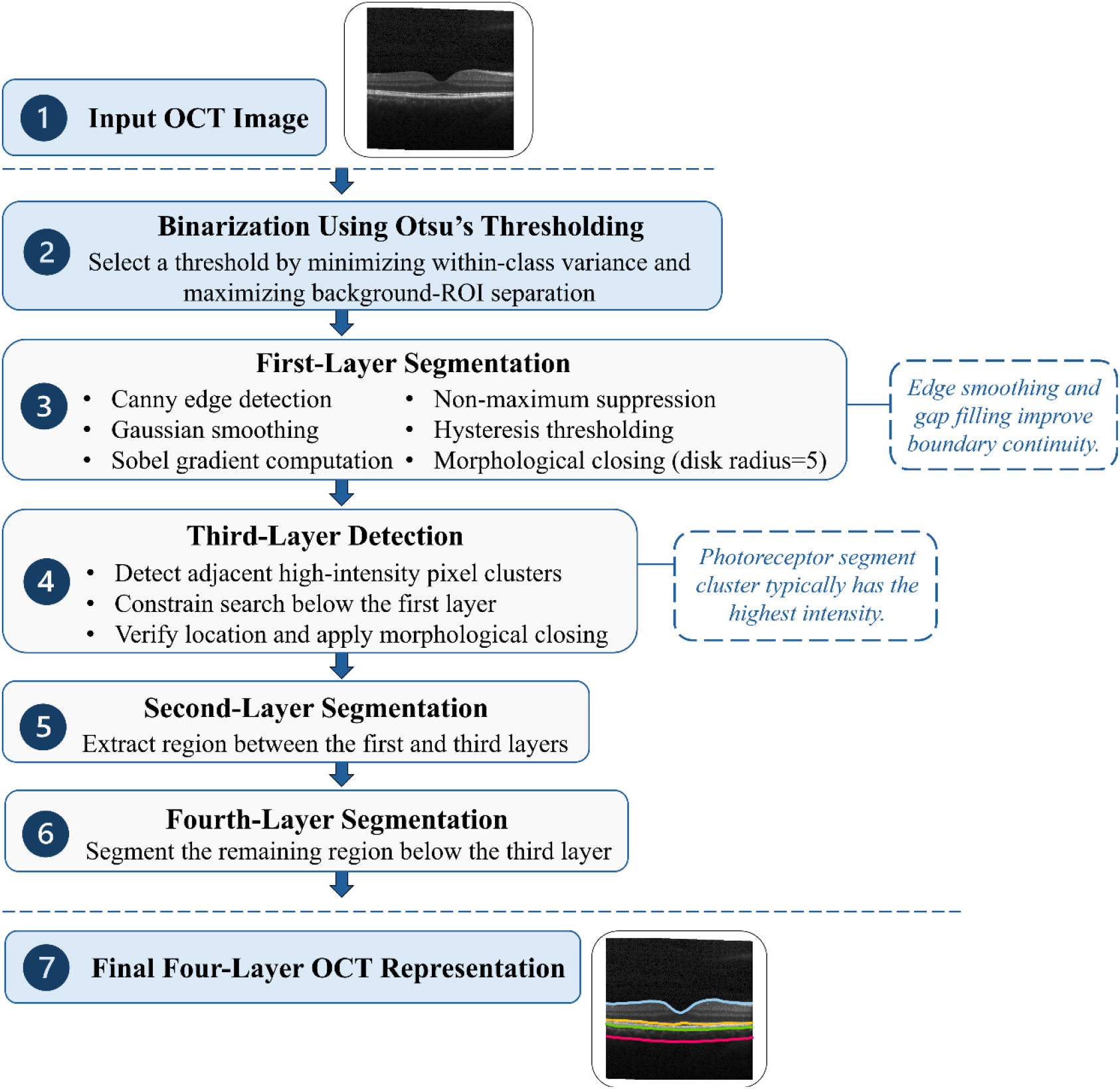
Schematic workflow of the proposed four-layer retinal OCT segmentation method.

### First and Third Layer Segmentation

The first layer’s (L1) segmentation includes Otsu’s method, edge detection, and morphological closing. It maximizes the variance between the background and the region of interest (ROI), ensuring that the threshold is selected to make the pixels in each region as homogeneous as possible. The method computes the weighted sum of variances across all possible threshold values and selects the threshold that minimizes within-class variance, effectively distinguishing the ROI from the background without requiring user-defined parameters. Canny edge detection was applied to the binarized images. In a multi-stage algorithm, the Canny filter first applies a Gaussian filter to the image to reduce noise, ensuring that minor variations in pixel intensity do not interfere with edge detection. Next, the algorithm computes the image’s intensity gradients by applying Sobel filters in both the horizontal and vertical directions, highlighting areas of rapid intensity change that may indicate edges. Non-maximum suppression is applied to sharpen these edges, keeping only the local maxima in the gradient direction and thinning the edges. Then, two thresholds (high and low) are applied using hysteresis thresholding. Pixels with gradient values above the high threshold are marked as edges, while those between the two thresholds are only considered edges if they are connected to a pixel above the high threshold. Morphological closing was used to smooth edges and close small gaps, holes, or breaks within an edge. It consists of two fundamental operations in which dilation is followed by erosion. Combining these operations improves the retention of the object’s overall size and shape while smoothing irregularities. The “disk’ structuring element with a radius of 5 is used. Figure 6a shows the L1 segmentation results.

**Figure 6.**
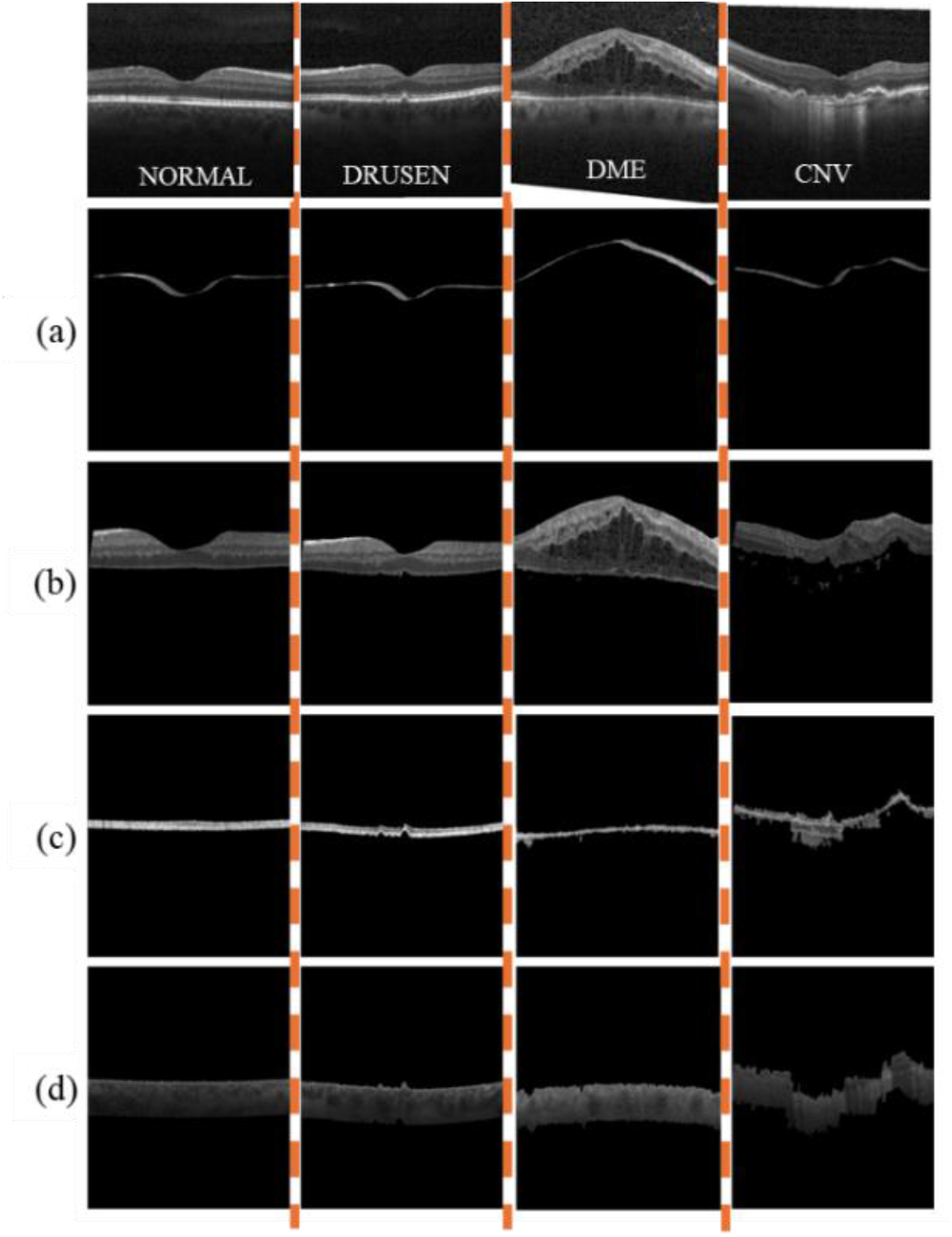
Examples of segmented retinal layers for Normal, Drusen, DME, and CNV OCT images illustrating the progression from raw input to anatomically separated retinal regions: original images, (a) first-layer segmentation, (b) second-layer, (c) third-layer segmentation, and (d) fourth-layer segmentation (Leyba-Mesa et al., 2025).

After segmentation of the L1, the third layer (L3) was identified. Clusters of adjacent pixels were detected from the binarized image. In this study, the photoreceptor segments correspond to the L3 and typically exhibit the highest intensity values in OCT images. As a result, the associated pixel cluster is often larger than the other clusters and is located below the first segmented layer. The proposed approach uses these characteristics to guide third-layer detection. However, the algorithm does not simply assume that the largest cluster corresponds to the third layer; instead, it uses the position of the segmented L1 to constrain and verify the L3’s location. A morphological closing operation was then applied to fill small gaps and connect discontinuities within the detected cluster. Figure 6c shows the resulting third-layer segmentation.

### Second and Fourth Layer Segmentation

The second layer (L2) lies between the first and third layers. The binary representations of these layers were used as boundaries for the second layer. Figure 6b shows the second-layer segmentation results for different classes. A remaining region under the third layer (L3) was segmented as the fourth layer. The approach enabled the extraction of the entire retinal structure, including the space beneath the retinal layers, ensuring a complete representation of the OCT image. Figure 6d shows the segmentation results of the fourth layer.

## Experimental Results and Discussion

### Feature Extraction from the Four Segmented Layers

We evaluated the contribution of the four segmented retinal layers to the classification of OCT diagnostic categories. A set of quantitative features was extracted from each layer, and one-way ANOVA was used to assess between-class differences and identify discriminative features. Because structural abnormalities associated with AMD and DME may appear as discontinuities, boundary irregularities, thickening, or fluid-filled regions, both statistical and texture-based descriptors were extracted. Low-order statistical moments, including mean, variance, skewness, and kurtosis, were computed from the segmented layers and the first derivative of the upper-layer boundaries to capture the intensity distribution and boundary variation. In DME cases, the third layer may show thickening or fluid pockets; therefore, texture features derived from the gray-level co-occurrence matrix (GLCM) (Park and Guldmann, 2020), including contrast, correlation, energy, and homogeneity, were used to characterize spatial intensity patterns within this layer. Contrast measures local intensity variation and highlights structural changes. Correlation reflects the linear dependency between neighboring gray levels and may help capture organized fluid-pocket patterns. Energy measures textural uniformity and indicates the presence of repetitive or dominant patterns, while homogeneity reflects the smoothness and uniformity of the texture. Before statistical analysis, Z-score standardization was applied to all features to ensure a common scale. The standardized value (Z) was calculated using Equation (5), *x* is the original feature value, *µ* is the mean value, and *σ* is the standard deviation. This transformation produces standardized features with a mean of 0 and a standard deviation of 1.

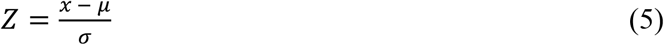

### Classification

Four supervised machine learning models, including K-Nearest Neighbors (KNN), Support Vector Machine (SVM), Naïve Bayes (NB), and Neural Network (NN), were implemented to evaluate the ability of features extracted from the segmented retinal layers to differentiate the four OCT categories. The model hyperparameters are the following: KNN used a cubic kernel with 10 neighbors, Minkowski distance with cubic weighting, while SVM also used a cubic kernel with automatic kernel scaling, a box constraint level of 1, and one-vs-one multiclass coding. Naïve Bayes was configured as a Gaussian Naïve Bayes model, and the neural network was a medium-sized network with one fully connected layer of 25 units and ReLU activation. The overall classification performance is presented in Table 1. The classifiers showed strong overall performance, suggesting that the segmented retinal layers provide discriminative structural information for OCT image classification. Among the four models, Naïve Bayes produced the lowest performance, which may be attributed to its assumption of feature independence. In this study, many of the extracted layer-based features are likely interrelated; therefore, this assumption may have reduced the effectiveness of the NB classifier. In contrast, the NN model achieved the highest overall performance, with an accuracy of 0.9817, sensitivity of 0.9788, specificity of 0.9938, and AUC of 0.9985. The trained neural network converged after 1000 iterations, with a final training loss of 0.013 and a step size of 0.01. The performance of the proposed NN-based classification approach was compared with previous studies, as shown in Table 2. The proposed approach achieved higher accuracy, sensitivity, and AUC than the compared methods, while maintaining high specificity. These results indicate that the proposed segmentation-based feature-extraction and classification framework is effective at distinguishing retinal OCT diagnostic categories.

**Table 1.** KNN, SVM, Naïve Bayes, and NN overall performance metrics (Leyba-Mesa et al., 2025).

|  | *Acc* | *Se* | *Sp* | *AUC* |
| --- | --- | --- | --- | --- |
| KNN | 0.8702 | 0.8236 | 0.9510 | 0.9708 |
| SVM | 0.9787 | 0.9748 | 0.9927 | 0.9948 |
| NB | 0.7286 | 0.6777 | 0.9005 | 0.8759 |
| NN | **0.9817** | **0.9788** | **0.9938** | **0.9985** |

**Table 2.** Comparison of the proposed model to previous works.

|  | *Acc* | *Se* | *Sp* | *AUC* |
| --- | --- | --- | --- | --- |
| Das et al. (2019) | 0.9550 | 0.9465 | 0.9854 | - |
| Hussain et al. (2018) | 0.9733 | 0.9467 | 1.0000 | 0.9733 |
| Rong et al. (2019) | 0.9286 | 0.9237 | 0.9340 | 0.9783 |
| Ai et al., (2022) | 0.987 | 0.987 | 0.996 | 0.991 |
| Sinha et al. (2023) | 0.944 | 0.944 | 0.9815 | 0.9983 |
| Huang (2023) | 0.978 | 0.978 | 0.9927 | 0.9983 |
| Khan (2025) | 0.975 | 0.9723 | N/A | N/A |
| Yang et al. (2025) | 0.9077 | 0.9262 | 0.9753 | 0.9809 |
| Leyba-Mesa et al. (2026) | 0.9758 | 0.9675 | 0.9913 | 0.9978 |
| **This work** | **0.9817** | **0.9788** | **0.9938** | **0.9985** |

### Feature Analysis and Ranking

The feature correlation heatmap (Figure 7) and the feature ranking results (Figure 8) were analyzed. This analysis provides insight into whether the selected features are redundant and which retinal layers contribute the most discriminative information for classifying diagnostic retinal OCT categories: Normal, CNV, DME, and Drusen.

**Figure 7.**
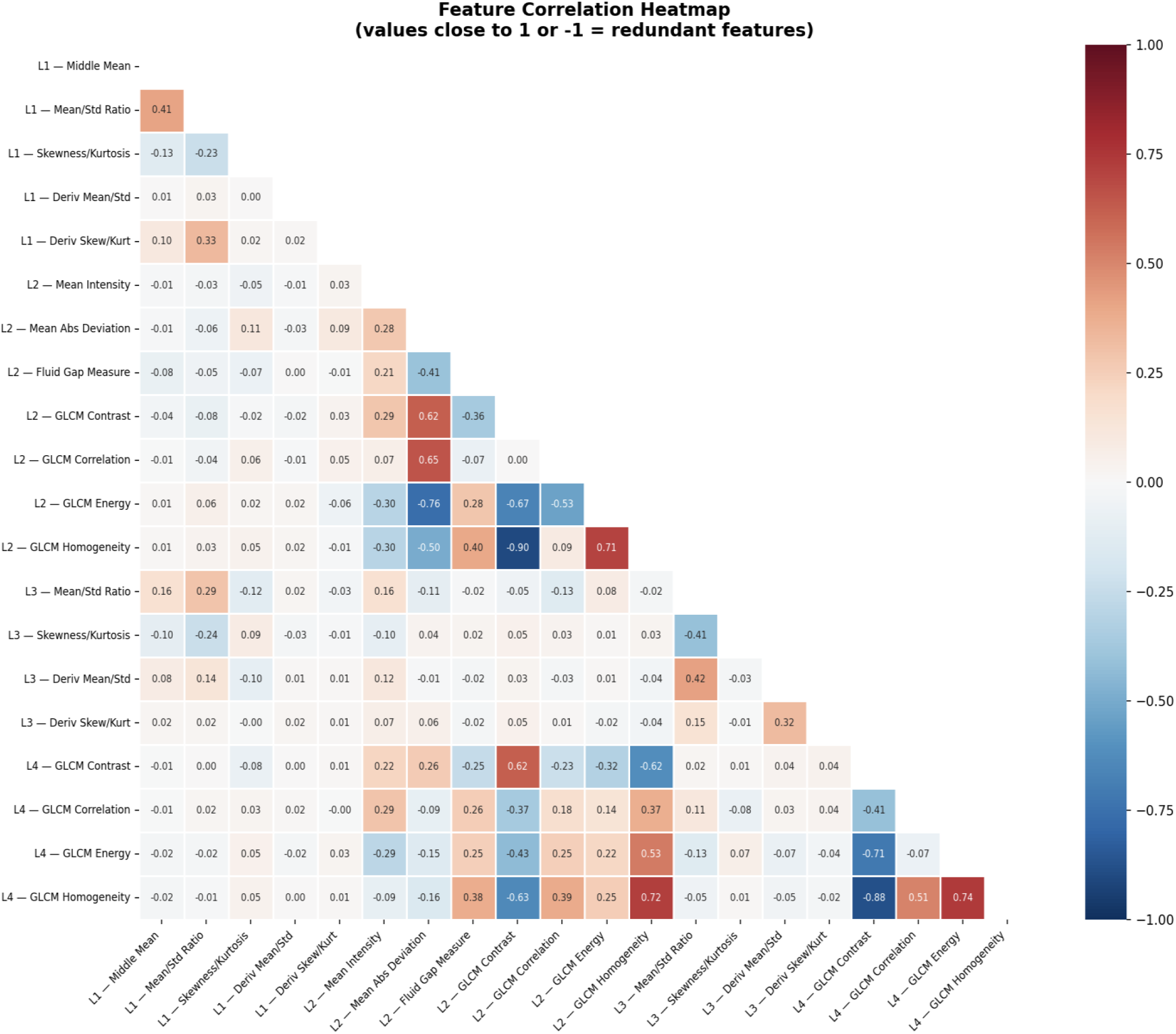
Feature correlation heatmap showing pairwise relationships among the extracted layer-based descriptors; values near 1 or -1 indicate strong redundancy, while weaker correlations indicate complementary features.

**Figure 8.**
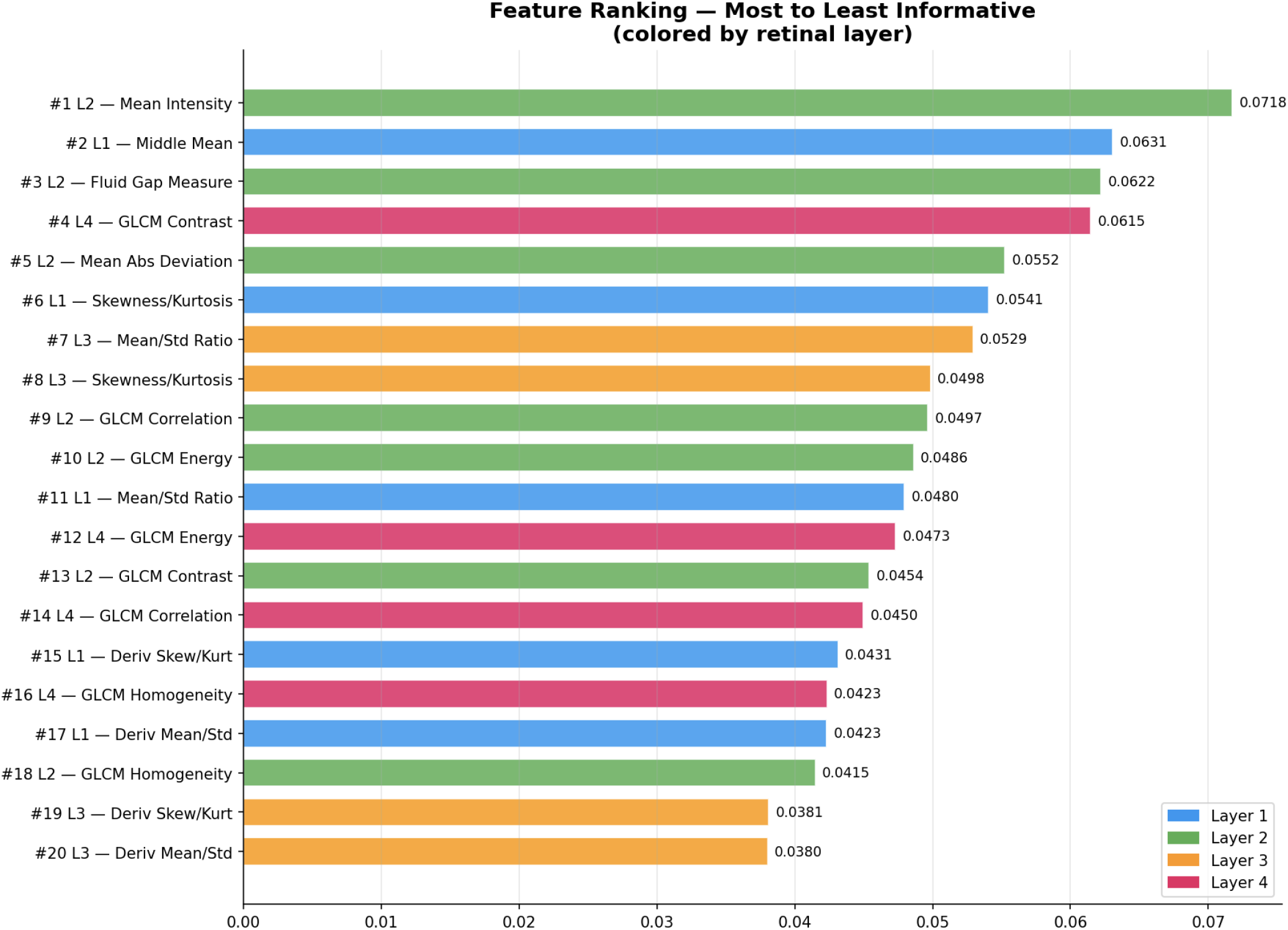
Feature ranking results for the extracted OCT descriptors, ordered from most to least informative and colored by retinal layer, highlighting each layer’s relative contribution to classification performance.

The feature correlation heatmap in Figure 7 shows that most pairwise relationships are weak to moderate, indicating that the proposed feature set is not dominated by severe multicollinearity. This is an important finding because the extracted descriptors were designed to represent distinct retinal characteristics, including intensity distribution, boundary morphology, derivative behavior, and intralayer texture. The absence of widespread near-perfect positive or negative correlations suggests that most features capture complementary rather than duplicate information, supporting their joint use in the classification framework. The strongest relationships are found among the GLCM-based texture features in L2 and L4. In these layers, contrast, correlation, energy, and homogeneity show moderate-to-strong associations, reflecting their shared role in characterizing spatial gray-level organization. Several of these relationships are negative, particularly between energy and contrast and between contrast and homogeneity, showing that as retinal texture becomes more irregular and heterogeneous, uniformity-related descriptors decrease while variation-related descriptors increase. This behavior aligns with pathological OCT patterns, in which disease disrupts the smooth layered appearance of the retina and produces irregular reflectivity distributions.

L2 showed strong internal structure in both correlation and ranking analyses. Many of its descriptors, mean intensity, mean absolute deviation, fluid gap measure, and GLCM features, were highly ranked. The top-ranked feature was L2 mean intensity, followed closely by L2 fluid gap measure and L2 mean absolute deviation. These results confirm that the L2 captures prominent disease-associated changes in reflectivity and local structural disruption. Because the L2 lies between the upper retinal boundary and the photoreceptor-associated region, this finding suggests that it is highly sensitive to abnormalities such as retinal thickening, fluid accumulation, and altered tissue organization. L4 also contributed to the classification of retinal OCT categories, particularly through texture descriptors. L4 GLCM contrast ranks fourth overall, and L4 GLCM energy, correlation, and homogeneity also ranked high (Figure 8).

In the heatmap, these features show moderate-to-strong mutual dependence. They collectively describe a coherent pattern of variation in subretinal and choroidal texture. Their prominence in the feature ranking suggests that the outer retinal and subretinal regions contain diagnostically important textural changes that help classify pathological cases from normal cases.

The statistical descriptors extracted from L1 and L3 generally showed lower pairwise correlations with the GLCM features and less internal dependence. The boundary- and derivative-based measurements captured information that differed structurally from that of the intralayer texture descriptors. For L1, features such as middle mean, skewness-to-kurtosis, mean-to-standard deviation ratio, and derivative statistics remain useful, with the L1 middle mean ranked second overall and several other L1 features appearing in the upper half of the ranking. These results suggest that the upper retinal boundary retains useful morphological information related to class differences, even if it is not the single dominant source of discriminative power.

Features from the L3 showed a distinct pattern. While the L3 mean-to-standard deviation ratio and the L3 skewness-to-kurtosis remain moderately important, the derivative-based L3 descriptors rank lowest in the ranking list. This may suggest that although the third layer contains relevant structural information, first-derivative statistics extracted from this region are less discriminative than the intensity and texture descriptors from L2 and L4. A likely interpretation is that L3 abnormalities are better captured by broader statistical characterization than by derivative-only measures, especially when compared with the more explicit fluid-related and textural signatures present in neighboring layers.

When the heatmap and ranking figure are interpreted together, a clear pattern emerges. The correlation heatmap confirms that the features are sufficiently diverse, while the ranking results identify which features carry the greatest class-separating value. The most informative descriptors are not confined to a single layer but are concentrated mainly in L2 and L4, with additional support from L1 and a more limited contribution from derivative features in L3. This layered distribution of importance supports the anatomical design of the segmentation framework, showing that disease-related information is spread across multiple retinal regions rather than being isolated to a single boundary or texture map. Overall, the low-to-moderate correlations across most feature pairs indicate that the extracted descriptors preserve complementary information, while the ranking results show that intensity-, fluid-, and texture-related measurements provide the strongest discrimination among OCT categories.

### Computational Costs Analysis

Computational usage was analyzed for the NN classifier, which was selected as the final model for the proposed OCT classification framework. We evaluated the NN classifier for runtime efficiency, model compactness, and training resource requirements. The results showed a prediction speed of approximately 160000 observations per second, a total training time of 2216.3 s, a compact model size of approximately 11 kB, and a coder model size of approximately 7 kB. Its high prediction speed and small model size support rapid processing and deployment in resource-constrained or high-throughput settings. The model achieved a 1.9% error rate and a validation cost of 1063, indicating stable training. Because classification is performed on extracted layer-specific features rather than raw OCT images, the framework remains lightweight, interpretable, and clinically relevant. This is an important advantage of the proposed framework, as it separates the clinically interpretable feature extraction stage from a compact and efficient classification stage.

## Conclusion

This study presented an adaptive and interpretable retinal OCT image analysis framework that combines preprocessing, anatomically guided four-layer retinal segmentation, layer-specific feature extraction, and machine-learning-based classification for Normal, CNV, DME, and Drusen cases. By operating on segmented retinal regions rather than the entire image, the proposed approach was designed to preserve clinically meaningful structure while improving transparency in the decision-making process. The experimental results demonstrated that the extracted layer-based descriptors provide a strong representation of retinal pathology across the four classes. Among the evaluated classifiers, the NN achieved the best overall performance, with an accuracy of 0.9817, sensitivity of 0.9788, specificity of 0.9938, and AUC of 0.9985, outperforming other fundamental models as well as some of the prior works included in the comparative analysis. Its performance was compatible with deep learning models. Feature analysis showed that complementary descriptors across multiple retinal layers contributed to classification, with L2 intensity- and fluid-related features and L4 texture features providing the strongest discriminative information. The final neural network combined high prediction speed with a very small memory footprint, supporting efficient deployment in automated OCT screening and clinical decision-support applications. The proposed work achieved high classification performance, offered insight into which retinal regions and features drive discrimination, and maintained low computational cost, making it a promising direction for transparent and practical OCT-based retinal disease analysis.

## Statements and Declarations

### Ethical considerations

This article does not contain any studies with human or animal participants.

### Data availability

N/A

## Acknowledgments

We acknowledge the creators and curators of the retinal OCT image dataset used in this work for making the data publicly available and enabling this research.

